# Methods for *in situ* quantitative root biology

**DOI:** 10.1101/2021.06.24.449808

**Authors:** Taras Pasternak, José Manuel Pérez-Pérez

**Affiliations:** Institute of Biology II/Molecular Plant Physiology, Centre for BioSystems Analysis, BIOSS Centre for Biological Signalling Studies University of Freiburg, 79104 Freiburg, Germany; Instituto de Bioingeniería, Universidad Miguel Hernández, 03202 Elche, Spain

**Author notes:** Corresponding authors: Taras Pasternak and José Manuel Pérez-Pérez. **One-sentence Summary:** Using the Deep Resolution Plant Phenotyping Platform, an epigenetic, cell cycle and geometrical atlas at subcellular resolution have been constructed for Arabidopsis, alfalfa and foxtail millet roots.

**Keywords:** root apical meristem (RAM), distal meristem, lateral root, cell cycle, chromatin structure, quantitative root biology *in situ*

## Abstract

When dealing with plant roots, a multi-scale description of the functional root structure is needed. Since the beginning of XXI century, new devices like laser confocal microscopes have been accessible for coarse root structure measurements, including 3D reconstruction. Most researchers are familiar with using simple 2D geometry visualization that does not allow quantitatively determination of key morphological features from an organ-like perspective. We provide here a detailed description of the quantitative methods available for three-dimensional (3D) analysis of root features at single cell resolution, including root asymmetry, lateral root analysis, xylem and phloem structure, cell cycle kinetics, and chromatin determination. Quantitative maps of the distal and proximal root meristems are shown for different species, including *Arabidopsis thaliana*, *Nicotiana tabacum* and *Medicago sativa*. A 3D analysis of the primary root tip showed divergence in chromatin organization and cell volume distribution between cell types and precisely mapped root zonation for each cell file. Detailed protocols are also provided. Possible pitfalls in the usage of the marker lines are discussed. Therefore, researchers who need to improve their quantitative root biology portfolio can use them as a reference.

## INTRODUCTION

The root is an essential organ required for plant anchor within the soil/substrate, and water and nutrients uptake. The root is a rapidly growing organ (up to 15 mm/per day in *Arabidopsis thaliana*, hereafter Arabidopsis), it is composed of a small distal meristem (columella and root cap) which protects the growing tip from mechanical cell damage during soil penetration, and a large proximal meristem responsible for overall growth (Bennett and Scheres 2010). The distal meristem also acts as a signaling hub for growth coordination in response to the soil environment (Barlow 2002).

Detailed investigation of the root structure was initiated more than 120 years ago, shortly after the building of the first optical microscopes. A detailed radial root structure with all cell files origin was thoroughly described at that time in Ranunculaceae (Maxwell 1893). Later on, with the development of DNA labeling techniques, timing and spatial location of mitotic cells were investigated in *Allium cepa* L. root meristems (Merriman 1904; Barsony 1927). Following these earlier studies, cell cycle kinetics in the root meristem was studied in pea (*Pisum sativum* L.) by quantifying the ratio of cells in a particular stage of the cell cycle to the total number of proliferating cells (Brown 1951). With the development of the thymidine incorporation assay (Das et al. 1957), more data about cell cycle kinetics in the root system became available. Namely, Clowes (1961) and Van’t Hof and Sparrow (1963) proposed a method based on H3-thymidine incorporation into the replicating DNA. Van’t Hof (1967) further modified this method to accumulate cells in mitosis by additional colchicine treatment. Nowadays, 5-ethynyl-20-deoxyuridine (EdU) incorporation has been used to study cell cycle kinetics in the root (Buck et al. 2008; Pasternak et al. 2021a). A detailed study of the primary root (PR) meristem of the Arabidopsis plant model was published by Dolan et al. (1993) one century after the pioneer work of Maxwell (1893).

Meristem zonation is another open question in plant root biology. It was earlier stated that meristem consists of cells with different cell cycle kinetics and different cell fates (Clowes 1961). However, root meristem size in Arabidopsis is usually defined based on the number of non-elongating cortex cells from 2D images, without any association to their proliferation activity and without considering the different tissues (Dello Ioio et al. 2007). Recently, Salvi et al. (2020) defined also the transition zone within the root meristem. However, it is not clear which cortex cells should be used to characterize the root meristem in other species like wheat, tomato, alfalfa, etc., which contain up to five cortex layers. Here we provide methods for precise connection of the cell size (volume) and cell proliferation activity in the root meristem. The next gap in root biology is the characterization of the so-called distal meristem (Barlow 2002; Bennett and Scheres 2010). In addition to its protective and signaling role, the distal meristem may serve as an ideal tool to study all cell development stages from cell division to terminal chromatin condensation, including cell death.

The kinetics of the cell cycle and its organ-wide distribution also has not brought enough attention by existing methods. Earlier work (Phillips and Torrey 1972) demonstrated that cell cycle duration is different in different cell files. Thus, the widely used CyclinB1;1 marker is not adequately reflecting all cells at the G2/M transition because their expression is dependent, among other factors, on the G2 duration in each cell, which is longer in the epidermis and the cortex (Lavrekha et al. 2017). To fill this gap, we described here a marker-free method that allows us to detect cell cycle and endocycle kinetics in all cell files at once.

Another missing feature in root biology is the detailed investigation of chromatin structure/organization *in situ*. Nuclei morphology is intrinsically linked to biological processes like gene expression regulation, cell fate establishment and cell differentiation (Conklin, 1912). Moreover, chromatin structure can also be used as a marker of cell cycle (Sequeira-Mendes and Gutierrez (2015). However, a robust and simple method for the analysis of a large number of nuclei within the root meristem and its link with cell fate and cell cycle kinetics are missing. As shown already by Maxwell (1893) and Dolan et al. (1993), QC cells have a very compact nucleus with a long (> 24 h) G1 phase. Zhang et al. (2013) claimed that cytokinin could induce QC cell division, but no additional evidence has been presented so far. For clarification of this point, a detailed investigation of the cell cycle kinetics in the QC region and quantification of the chromatin status at spatial resolution is required.

The proposed protocol will allow performing such analyses and connect nuclei landscape with cell cycle stages and cell differentiation status at a whole-organ level. The protocol described here for *in situ* quantitative root biology can be applied to any plant species, including dicotyledonous and monocotyledonous plants, which largely differ in their root system architectures. In the current paper, we provide a comprehensive description of cell biology approaches to fulfill most of the existing gaps in root biology and to characterize the root system at a single cell resolution.

## MATERIALS AND METHODS

### Plant material and growth conditions

Seeds of *Arabidopsis thaliana*, *Nicotiana tabacum*, *Medicago sativa*, and *Setaria italica* and the growth conditions used have been thoroughly described elsewhere (Pasternak et al. 2015; 2017; Tang et al. 2021).

### Plant material preparation and labelling

All procedures for plant fixation, treatments, and imaging have been described previously (Pasternak et al. 2015; 2017; 2020). The main differences from the previously published protocols are presented below.

#### Cell boundary labeling

Our cell boundary labeling protocol is based on the binding of propidium iodide to de-ketonized cell wall polysaccharides at low pH (1.4) in the presence of sulfur. Although the basic protocol has been described previously (Truernit et al. 2008), significant modifications are required to adapt the protocol for 3D scanning and analysis of the inner tissue layers, including xylem and phloem tissues. One limitation is that the original fixation in acetic acid leads to significant tissue maceration and often damages the softer mature plant parts. For this reason, we recommend fixation with formaldehyde in MTSB buffer at pH 7.0 before labeling. The de-ketonization level (time of periodic acid treatment) is another crucial parameter. For Arabidopsis roots, a 30 min de-ketonization in 1% periodic acid partially punctured the cell walls, especially in the vasculature with thinner cell walls. We recommend reducing the de-ketonization time to 15–20 minutes. The mounting procedure is another crucial step. As already mentioned, the spacer should have a similar thickness to the object. A thick object can be scanned using double-sided scanning by mounting the samples between 2 coverslips: a 24 × 60 mm size as a base and a 24 × 32 mm size as a cover. This adjustment allows the object to be scanned from both sides to avoid a low signal-to-noise ratio in the deeper parts.

#### Chromatin and cell cycle detection

For detection of the chromatin and cell cycle, plants were transferred to a 12-well plate with appropriate liquid medium (TK1 for Arabidopsis) for 12 h. Thereafter, 10 μM 5-ethynyl-20-deoxyuridine (EdU) was added for appropriate incubation period. Seedlings were then fixed and subjected to detection EdU detection and scanning as described (Pasternak et al. 2015). For chromatin analysis, we highly recommend to scan images with voxel size 0.2 × 0.2 × 0.3 (X-Y-Z resolution). Dynamical corrections of the scanning also require keeping the signal intensity and signal-to-noise ratio as constant as possible through whole stacks. To reach this aim, one may adjust detector gain and the average scan numbers for each sample.

#### Preservation of the 3D structure

Preservation of the 3D structure is crucial for properly analyzing the cell and nucleus shape and cell position. The most sensitive issue to the shape disturbing is mature cells with large vacuoles that can quickly shrink under not mild conditions. We have used several tricks to prevent shape disturbing. We minimize the time of de-ketonization and cell wall digestions to keep cell wall thinner. However, the most crucial steps are methanol treatment and the mounting procedure. For the methanol, we suggest adding water to methanol very gradually but do not transfer samples to new solution. For the mounting, since almost all mounting mediums are glycerol-based, direct transfer to mounting medium after labeling eventually led to disturbing cell shape. To avoid this issue, we gradually add 50% glycerol to the samples before mounting them to the final concentration (25% glycerol) in the mounting medium. This trick allows keeping the original shape of the cells even in the differentiation zone of the root.

#### Image processing and analysis

Image analysis were performed essentially according to Pasternak et al. (2021). Briefly, images were converted to hdf5 format using the LOCI plugin for ImageJ (http://imagej.nih.gov/ij), then stitched together to obtain a root tip total length of 400 μm from the QC using xuvTools (Emmenlauer et al. 2009). Finally, 5-10 representative roots were chosen for detailed annotation. The DAPI and EdU channel images were processed with the iRoCS Toolbox (Schmidt et al. 2014) in the following way: nuclei were automatically detected using the “01-Detect Nuclei” plugin, then the epidermis was semi-automatically labelled using the “02-Label Epidermis” plugin. After the QC position was marked (Channel->New Annotation Channel), the nuclei were set in spherical coordinates using the “03-Attach iRoCS” plugin. Automatic classification of the nuclei to the corresponding cell types (root cap [RC], columella [col], epidermis [epi], cortex [co], endodermis [en], pericycle [pe], vasculature [va]) was done using the “04-Assign Layers” plugin, which also enabled the automatic annotation of nuclei in mitotic state (option “Re-classify mitotic state”). All annotated roots were manually corrected for erroneous layer, mitosis, and EdU assignments.

#### Nuclei segmentation and analysis

For the nucleus geometry analysis, original nuclei-labeled images were converted to .tiff, and OTZU methods were applied (Otsu 1979). Thereafter nucleus edges have been detected, images were converted to hdf5 and further segmentation was performed accordingly to the standard pipeline (Pasternak et al. 2021b). Individual nuclei analyses were performed according to Dubos et al. (2020) with slight modifications. Single layer images from a given cell type and area containing 20-30 nuclei have been selected, and processed with the NucleusJ2.0 plugin as described elsewhere (Dubos et al. 2020).

## RESULTS

Despite their simple structure, roots are complex organs and we can quantify their morphology at different biological scales, ranging from proteins, organelles, single cells and tissues. Based on these scales, we have divided the pipeline of analysis into four modules: (I) nuclei and chromatin analysis; (II) cell cycle analysis; (III) protein and protein complex localization; and (IV) cell and organ geometry analysis (Figure 1). We included an X-Y-Z coordinate system (Schmidt et al. 2014) that allows us to extract positional information for each cell within a single root. This approach was applied to study the root apical meristem (RAM) and the differentiation zone of the primary roots (PRs), as well as the young lateral roots (LRs), of several plant species. Moreover, the modules can be combined, thus the simultaneous extraction of cell geometry, cell cycle and chromatin organization are feasible.

**Figure 1.**
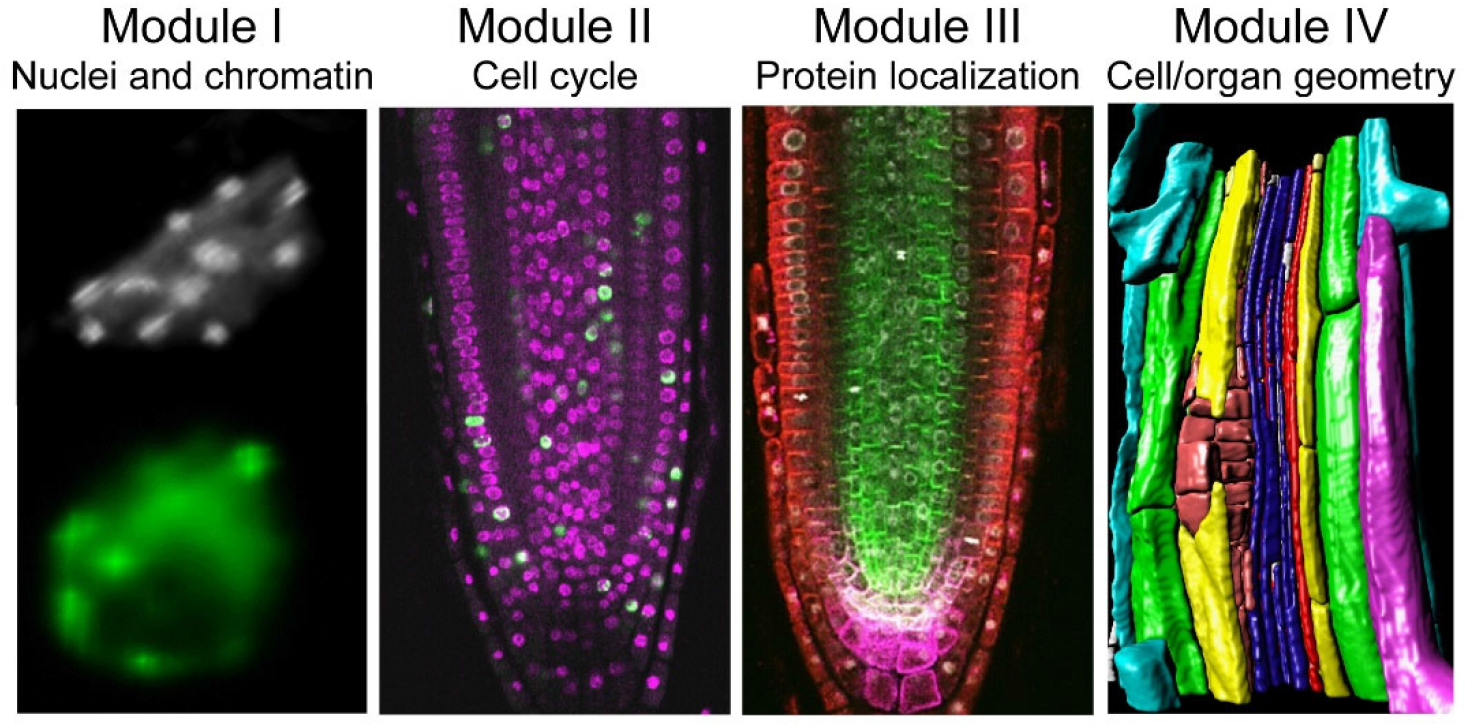
Proposed pipeline for *in situ* quantitative root biology. We defined four different modules for cellular phenotyping of serial confocal images. Each cell within a single root is defined by a 3D coordinate system (Schmidt et al. 2014) and the proposed method allow simultaneous extraction of cell geometry, cell cycle and chromatin features.

### Module I: Nuclei and chromatin analysis

The plant nucleus is not an inactive container of the DNA and its dynamic organization plays a crucial role in coordinating DNA replication, DNA transcription, and cell fate. The plant nucleus shows specific cell fate-dependent and position-dependent structural, compositional and functional features. Namely, specific characteristics of the nuclear shape, size, distribution and composition of nuclear domains, heterochromatin content and chromatin condensation serve as precise fingerprint of the cell fate and cell cycle status. These features allow dissecting all chromatin statuses, including the origin of DNA replication, histone modification, and ratio between hetero and euchromatin, among others. The aim of module I is to provide a detailed chromatin functional atlas of the root.

First, we investigated nuclei organization in Arabidopsis primary roots (PRs; Figure 2A). We observed substantial differences in nuclei morphology between cell types, especially between root cap (RC), cortex/epidermis and the more inner tissue layers (Figure 2B). To quantify these differences, we performed volumetric nuclei analysis. We found rapid divergence of the nuclei volume and nuclei heterogeneity between cell types even in the division zone of the RAM, with higher nuclei volume in the cortex and epidermis layers (Figure 2C). Next, we extended our analysis to single nuclei features by using the NucleusJ2.0 plugin that allowed us to extract up to 12 morphological parameters per nucleus (Dubos et al. 2020). When studying cells of the same tissue layer in the PR, we can distinguish different cell cycle stages in the proliferation zone of the RAM based on nuclei volume and other morphological features (Figure 2D, E).

**Figure 2.**
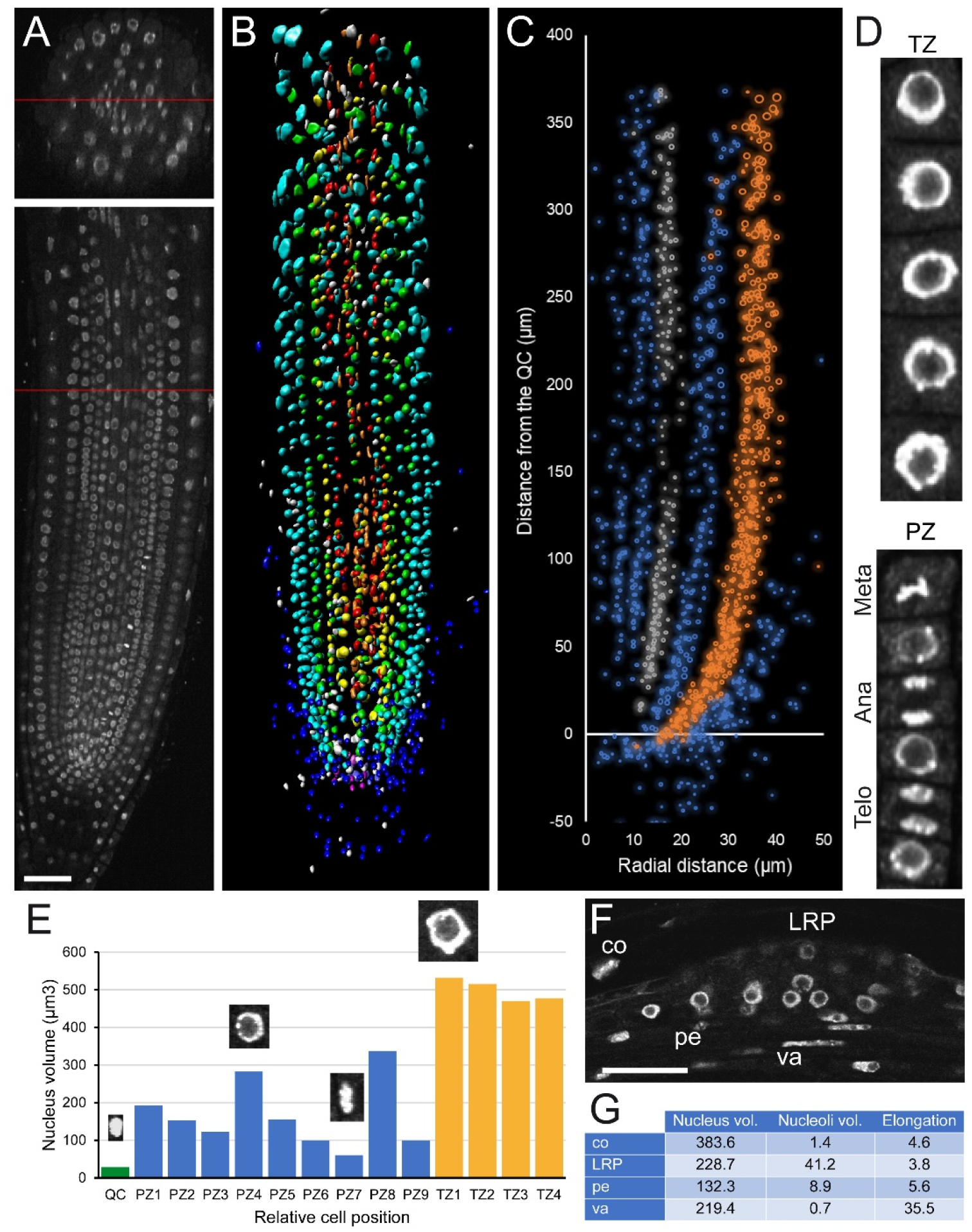
Analysis of the chromatin structure after propidium iodide labelling. (A) Confocal 3D image of an Arabidopsis PR; top: cross section, bottom: longitudinal section. (B) Overview of the nuclei volumetric analysis; each color represents nuclei from same tissue. (C) Nuclei distribution with radius as cell volume along the longitudinal and radial axes of different tissue layers; epidermis cells are depicted in orange and pericycle cells are shown in grey. (D) Selected nuclei of epidermal cells in the proliferation zone (PZ) and the transition zone (TZ) of the RAM for detailed analysis. Meta: metaphase, Ana: anaphase, Telo: telophase. (E) Nuclei volume of individual cells. QC: quiescent center. Some nuclei images are shown. (F) A representative 2D image after DAPI labeling in a newly formed LR primordium (LRP). Individual cells used for data extraction are shown (co: cortex, pe: pericycle, va: vascular). (G) Quantitative data from individual cells. Volumes are in μm^3^. Scale bars: 50 μm.

Besides the RAM, elongation and differentiation zones of the root have a key importance for unveiling the mechanism of LR formation and nodule/pseudo nodule induction. In the RAM, each cell has its own nuclei structure fingerprint, including nucleolus size, which varies according to its cell type and its position. By applying module I, some of these features have been measured on an early LR primordium arising from the PR in Arabidopsis (Figure 2F and Supplemental Figure S1), and a comparison with the nuclei features from the same tissue in PRs is also possible.

DAPI staining of nuclei for determining cell cycle stages and chromatin status in roots is a marker-free strategy that could be used in non-model plants. We demonstrated that such analysis is also possible for alfalfa (*Medicago sativa*), and we clearly distinguished different cell cycle stages (G1 and mitotic cells) based on nuclei volume and other morphological features (Figure 3A-E). Next, we applied module I analysis to tobacco (*Nicotiana tabacum* L.) roots. In contrast to that found in Arabidopsis, cortex cells in the mature region of the tobacco PR are able to divide and even induced the pseudo-nodules (Arora et al. 1959). Detailed chromatin structure analysis shown that cortex cells in tobacco contained regular nuclei structure with similar sphericity and nuclei volume as in the pericycle cells, although the later displayed higher nuclei occupancy (Figure 3F-I).

**Figure 3.**
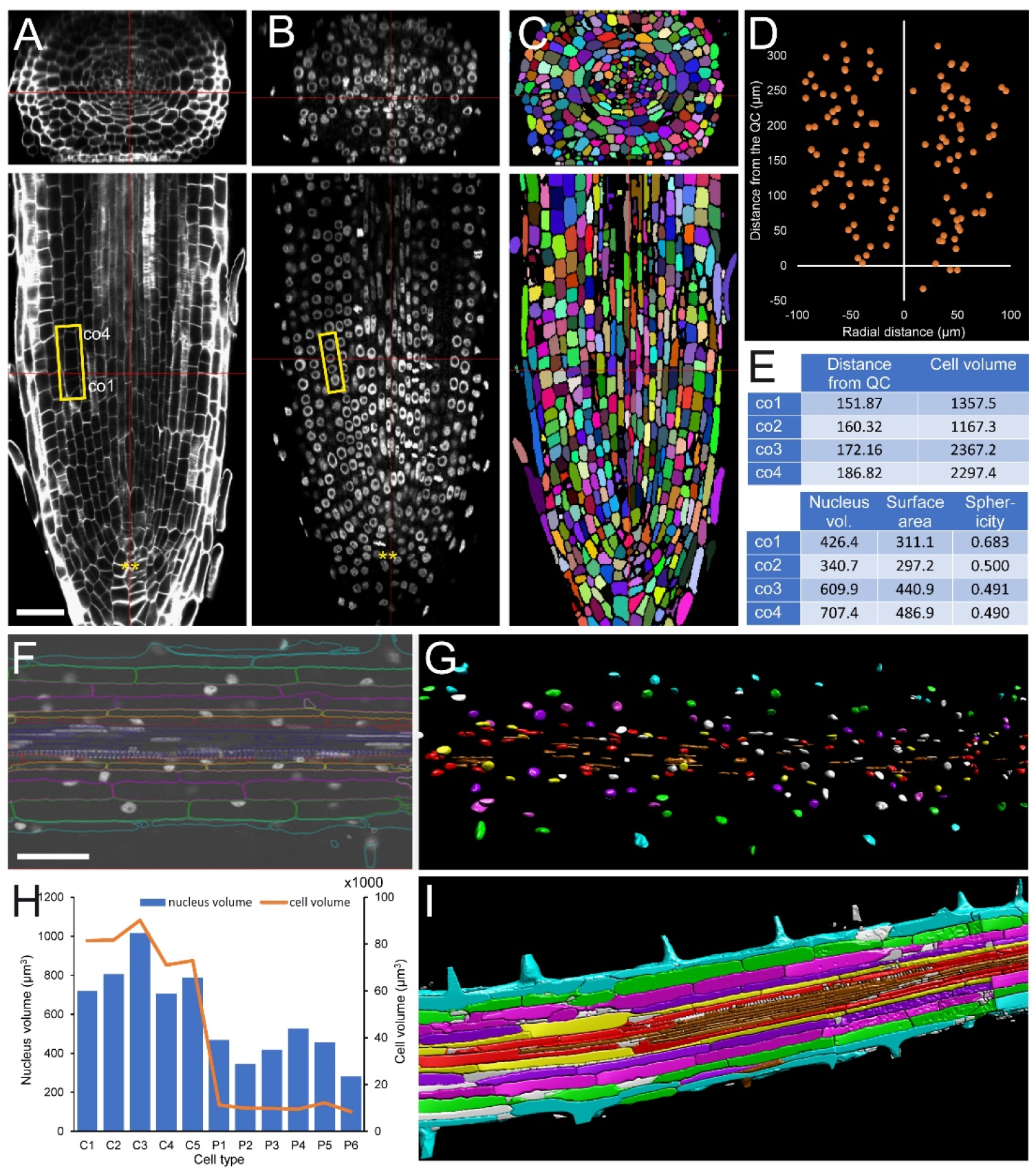
Nuclei and chromatin analysis in *Medicago sativa* and tobacco roots. Roots were fixed, double-labeled for cell border and nucleus, scanned and analyzed as described in Materials and Methods. (A) Cell wall and (B) nuclei labeling in the PR of *Medicago sativa*. Yellow rectangles in A, B: cells and nuclei chosen for detailed analysis. (C) Cell border labelling and cell segmentation. (D) Mitosis distribution along the longitudinal and radial axes in all root tissues. (E) Details of morphological features of selected cells and nuclei from A, B. Distances are in μm, areas in μm^2^, and volumes in μm^3^. (F) A representative 2D image in the mature region of a tobacco PR (G) Nuclei segmentation of the PR shown in F; different colors indicate nuclei from the same cell layers. (H) Nuclei volume and cell volume representation in individual cells; C: cortex, P: pericycle. (I) 3D render of the cell border labeling and segmentation. Each color represents individual cell layer. Scale bars:50 μm.

### Module II: Cell cycle analysis

Contributing new cells from the RAM is critical for root growth. Cell cycle progression is highly and dynamically regulated at the chromatin level. In module II, we combined two DNA staining and marker-free methods to identify different cell cycle stages (Figure 4A-C): (a) 5-ethynyl-2 deoxyuridine (EdU) staining that marks DNA replicative nuclei, and (b) nuclei labeling with DAPI that allows scoring mitotic figures. Thus, we can distinguish cells in four different statuses: (1) cells with very compact (but regular) chromatin that prevent nucleosome/chromatin disassemble (*i.e.* DNA replication), (2, 3) cells in G1 and G2 in the division zone of the meristem with more relaxed chromatin (Figure 4D), and (4) irregular chromatin in differentiation and mature zone. Incubation with EdU during variable times allows investigation of the kinetics of DNA replication, as a short EdU incubation marks only a small proportion of cells in the meristem (Figure 4E), while longer EdU incubation marks most cells in the proximal meristem but not the QC region (Supplemental Figure S2).

**Figure 4.**
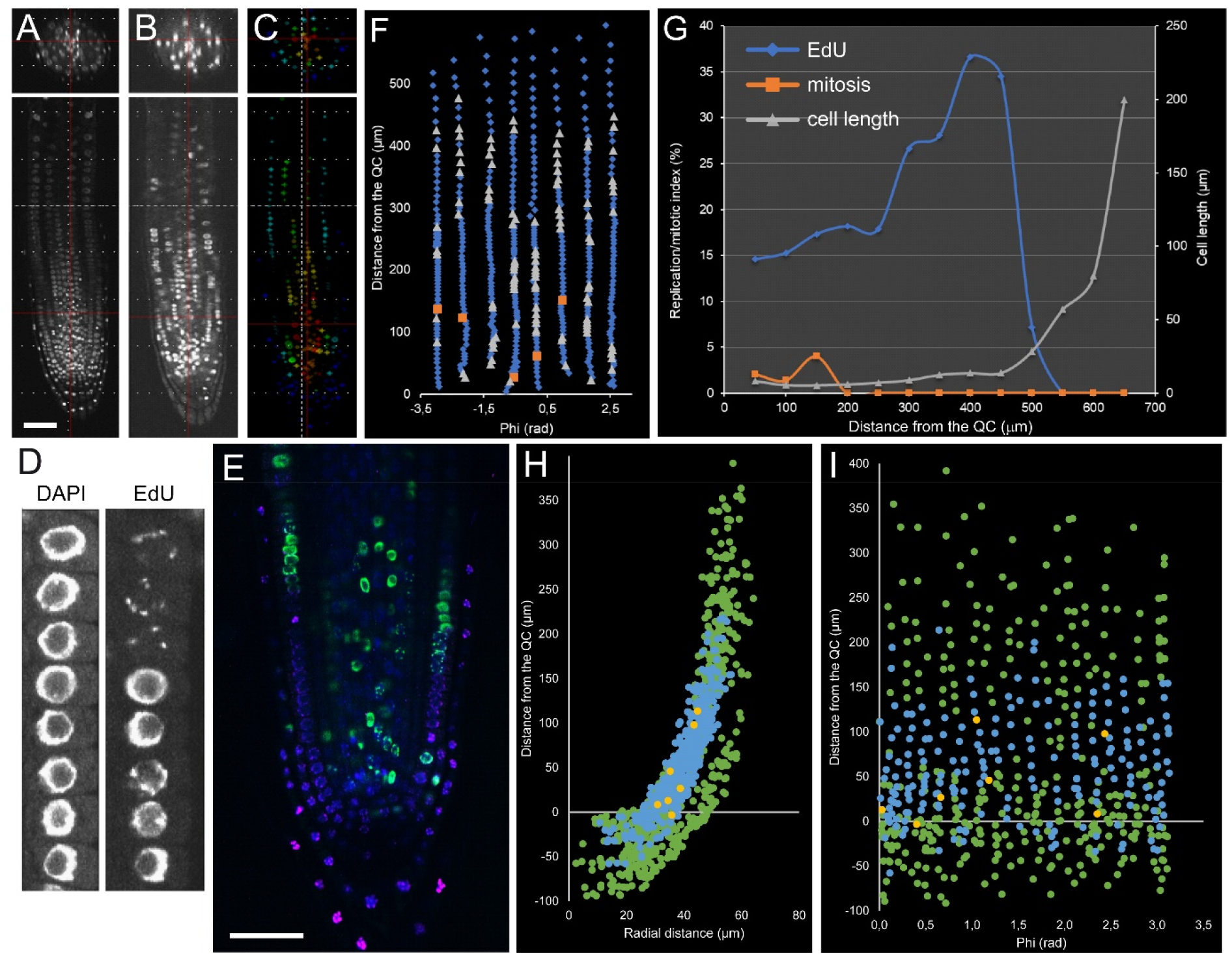
Detection of key cell cycle events in Arabidopsis PR. (A) DAPI labeling for nuclei and mitosis identification. (B) EdU staining (20 min) for detection of DNA replicative cells. (C) EdU-positive nuclei segmented by cell layer. (D) DAPI and EdU staining on cortex ceels in the RAM. (E) Unrolled layer of cortex tissue. Blue: interphase nuclei; grey: DNA replicative nuclei; orange: mitotic nuclei. (F) Quantitative analysis of DNA replication (blue) and mitotic index (orange) in the cortex tissue along the longitudinal axis. Cell lengths are in grey (G-I) DNA replication in the Arabidopsis distal meristem (DM). Seedlings were incubated with EdU for 90 min (G; DAPI: blue, EdU: green) and for 8 h (H; DAPI: green, EdU: blue). (I) Unrolled layers of RC. Green: all nuclei; blue: EdU-positive nuclei; yellow: mitotic nuclei. Scale bars: 50 μm.

Module II allowed us to study cell cycle within a single root at cellular resolution. For example, cortex tissue in the proliferation zone of the meristem contained 258 cells (located from 0-200 μm from the QC), 49 of which had EdU-positive nuclei, and 5 were in mitosis; while the transition zone (TZ) of the meristem (200-400 μm from the QC) contained 217 cells with 48 cells with EdU-positive nuclei and no mitotic cells (Figure 4F-G). Similar data can be extracted for all other tissue layers as well. This kind of analysis has been used previously for quantifying root zonation in each cell file, and led to the conclusion that the meristem size is tissue-layer specific (Pasternak et al. 2017).

Module II (in combination with Module I) analysis can also be used for stem cell niche studies. The stem cell niche is defined based on the QC cells, which are quiescent stem cells with very long G1 duration (Supplemental Figure S2). The long G1 duration of the QC might be dependent on their very dense nuclei (Figure 2E) that difficult DNA replication. Hence, module I and II analysis in the stem cell region using mutants and different environmental signals will provide additional insight into the biology of plant stem cells (Svolacchia et al. 2020).

Contrary to the proximal meristem, the distal meristem (composed of central columella and lateral root cap [RC] cells) is not involved directly in root growth but it plays an essential role in protecting the root against abrasive damage of the soil and it is involved in the perception of several environmental signals. The distal meristem is a tissue with rapid turnover of short-lived cells, which is regulated by an intricate balance of new cell production, differentiation, and programmed cell death (Xuan et al. 2016). The precise kinetics of distal meristem cell production and differentiation has not yet been studied. By combining module I and II, we found that the outer layers of the RC do not contribute to new cell production (Figure 4H-I).

Detailed analysis of cell cycle events in the differentiation zone of the root shows that almost all cell types replicate their DNA. However, only pericycle cells can fulfill mitosis in this region during LR formation in Arabidopsis. On the other hand, cortex nuclei in *Medicago sativa* and tobacco PRs keep regular structure layers (close to round nuclei with regular chromocenter distribution), which might account for pseudonodule induction in tobacco and nodule induction in legumes (Supplemental Figure S3).

Another unknown issue in root biology is how phytohormones regulate cell cycle transitions, which is key to understanding plant growth, tissue regeneration, and stress responses (Shimotono et al. 2021). The auxin effect on cell cycle regulation has not been studied at cellular resolution so far. Module II analysis will allow closing this gap. We found that short incubation with 100 nM of the synthetic auxin NAA significantly inhibited DNA replication and cell division of all cells within the proliferation region of the RAM, while it induced DNA replication of pericycle cells in the transition region of the RAM, which are the source of the new LRs (Figure 5).

**Figure 5.**
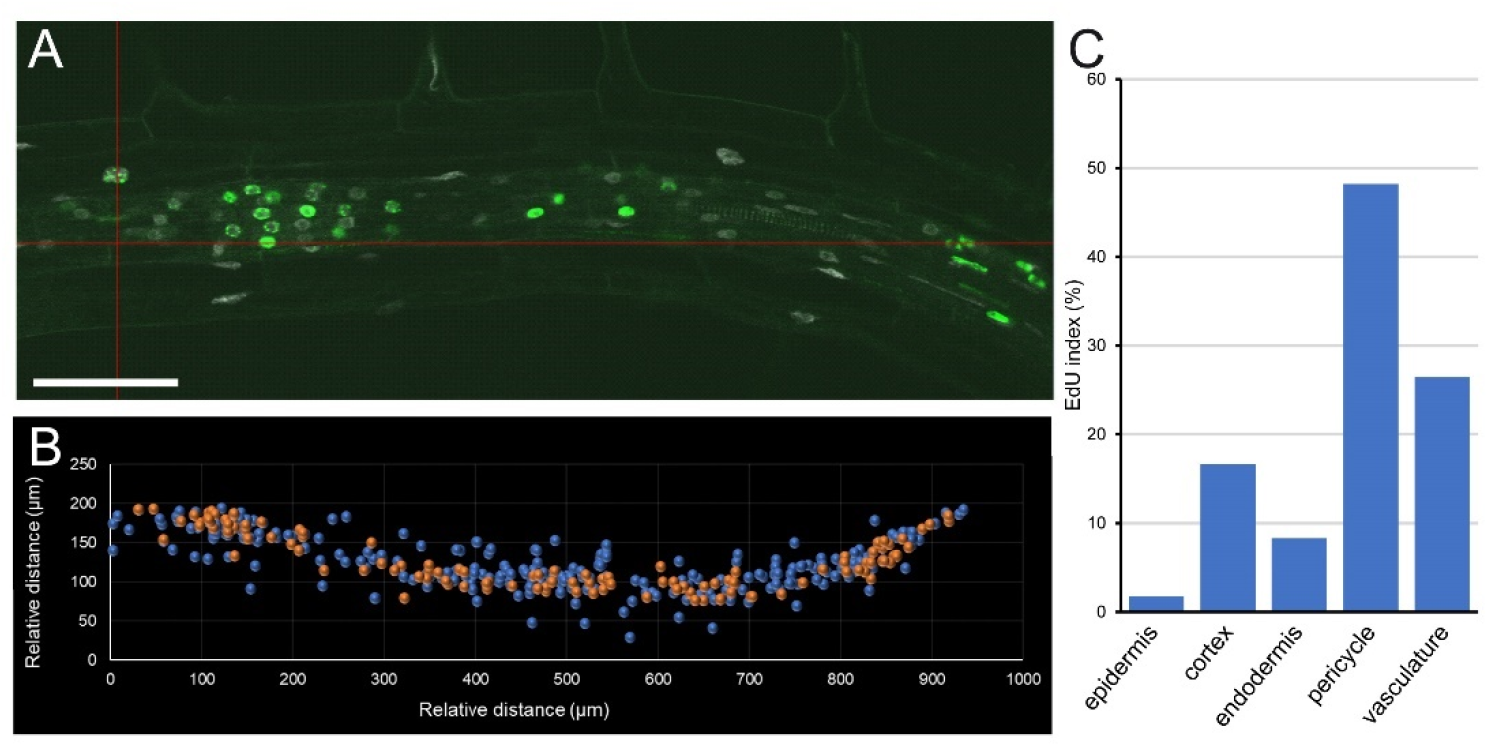
Effect of auxin on cell cycle events in Arabidopsis PR. Seedlings have been treated for 16 h with 100 nM NAA, EdU has been added for 90 minutes, samples have been fixed and analyzed as described in Materials and Methods. (A) Confocal 2D image; nuclei are in white; EdU are in green. (B) Position of the nucleus (blue) and EdU-positive nuclei (orange). (C) Cumulative EdU index in different cell types. Scale bar: 50 μm.

In conclusion, module II allows us to quantify cell cycle events *in situ* for all cell types and calculate the kinetics of the cell cycle (Pasternak et al. 2021a).

### Module III: Protein and protein complex localization

As inferred from the results of module I, each single cell within the root from the same or different tissues displayed a specific nucleus morphology, which would affect their gene expression and protein activity levels. This, in turn, made questionable classical molecular biology methods of protein level quantification and analysis (*i.e.* Western blot) and demands a robust method to determine protein localization and protein levels at single cell resolution.

Protein localization in plant cells is commonly studied using fluorescent protein fusions (Tanz et al. 2013). However, this approach requires obtaining stably transformed plants, which is time-consuming and it is limited to some plant species with available transformation and tissue culture protocols. In addition, the presence of a large hydrophilic region of the fluorescent protein may produce recombinant proteins with slightly different characteristics than the native proteins which, in turn, can affect their spatial localization.

Protein immunolocalization with a custom antibody against the protein of interest is a good alternative to study protein localization, as the antibodies can be assayed directly in different genetic backgrounds, including mutants, without the need of further crossing. If the antibody design is adequate, cross-reactivity in multiple plant species is possible (e.g. PIN1 and tubulin antibodies [Pasternak et al., 2015; Omelyanchuk et al. 2016], histone variant antibodies, etc.). One additional advantage of immunolocalization is the possibility of performing triple or quadruple labelling and studying protein-protein interaction *in situ*. Commonly used methods of studying a protein complex by pull-down or co-immunoprecipitation require cell lysis that might change protein conformation, hence they may not exactly allocate subcellular protein complexes. Moreover, protein complex composition may differ also between neighboring cells. Hence, *in situ* detection of a protein complex may provide adequate information about its functional activity.

Module III includes a detailed protocol for double or triple labelling with different antibodies as well as a proximity ligation assay (PLA assay) to study protein complex interactions (Pasternak et al., 2018). As an example of module III output, Figure 6A-B shows the immunolocalization of the auxin efflux facilitator membrane proteins PIN-FORMED1 (PIN1), PIN2 and PIN4 in the Arabidopsis RAM. Quantitative analysis of the PIN1-PIN4 complex localization and levels in 3D has been presented previously (Teale *et al.* 2021). Another example of module III application is the localization of different histone variants, due to most of the histone antibodies that are commercially available recognize conserved peptides of both plant and animal histones. Figure 6C-D shows an example of the H3K9me2 mark that serve as a key hub for regulation of chromatin status and gene expression (Zhang et al. 2018).

**Figure 6.**
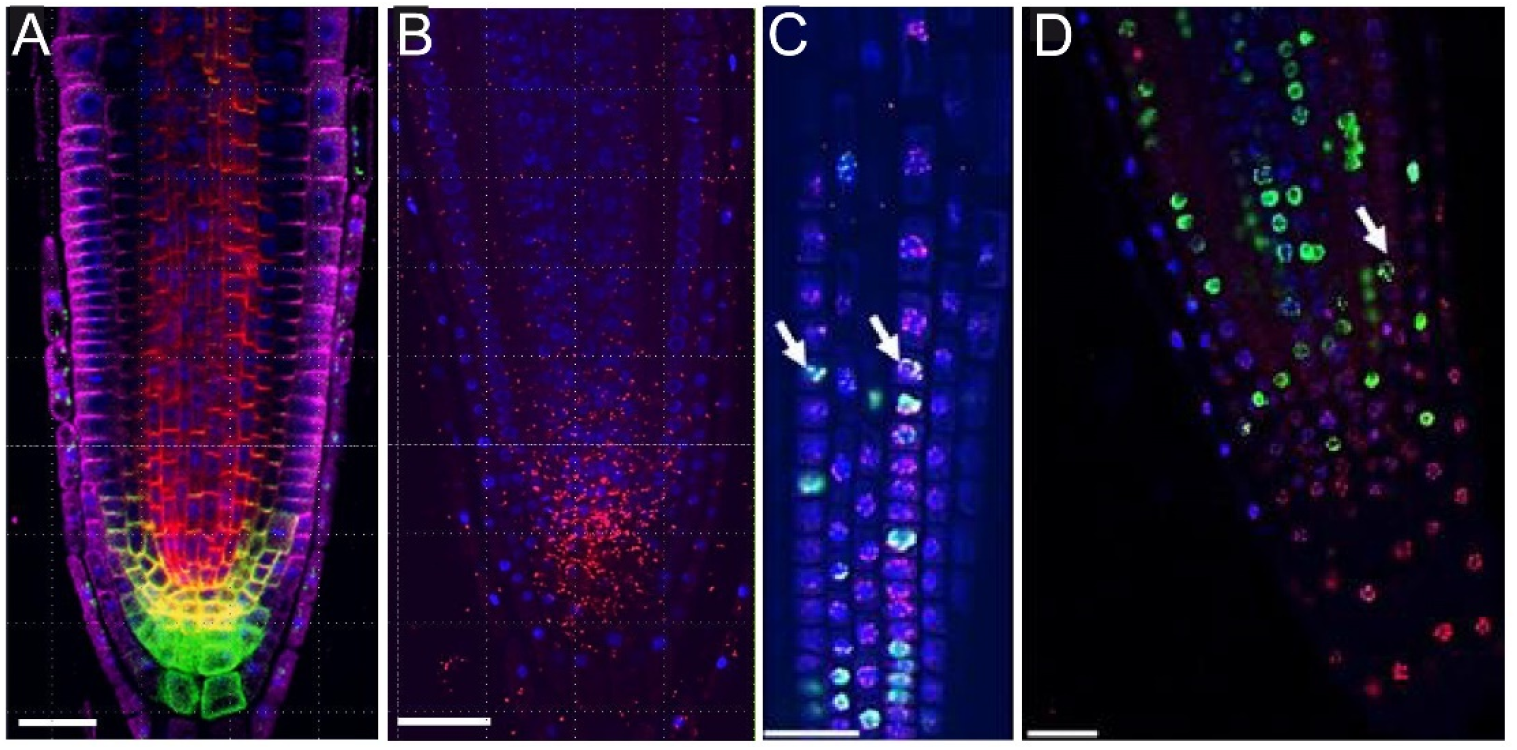
Protein and protein complex localization in Arabidopsis roots. (A-B) Detection of PIN1 and PIN4 complex by proximity ligation assay in Arabidopsis PR. (A) Immunolocalization: PIN1 (red), PIN2 (magenta) and PIN4 (green). (B) Proximity ligation assay: PIN1-PIN4 complex (red). Nuclei are indicated in blue. (C, D) Colocalization of the H3K9me2 with DNA replication events. DAPI is in blue; EdU (90 min incubation) in green and H3K9me2 immunolocalization is in magenta. Arrows marked visible colocalization of DNA replications with H3K9me2. Scale bars: 20 μm.

### Module IV: Cell and organ geometry analysis

For module IV analysis of Arabidopsis and *Setaria italica* PRs, individual cells were segmented and assigned to different tissue layers as described elsewhere (Pasternak *et. al.* 2021b; Figure 7A, D). In both species we found that cell volume was highly dependent on cell type. The outer tissue layers (epidermis and cortex) accounted for more than 80% of the total of the root volume (Figure 7B, E). Other cell types, such as those of the vasculature in the Arabidopsis RAM, are characterized by their small contribution to total root volume and large numbers (Figure 7B). Also, increase in cell volume of neighboring cells along the longitudinal axis was both cell type-specific and region-specific, with the higher values in epidermis and cortex cells (Figure 7C). Vasculature occupies the most central region of the root and it is mainly involved in both basipetal and acropetal transport (Figure 7F). Phloem is responsible for the transport of auxin and sucrose to the root, while xylem transports water and mineral nutrients to the shoots. In C4 plants, such as *Setaria italica*, cell volume of central metaxylem (MX) and protoxylem (PX) were extremely large, in opposing contrast to phloem cells (Supplemental Figure S5).

**Figure 7.**
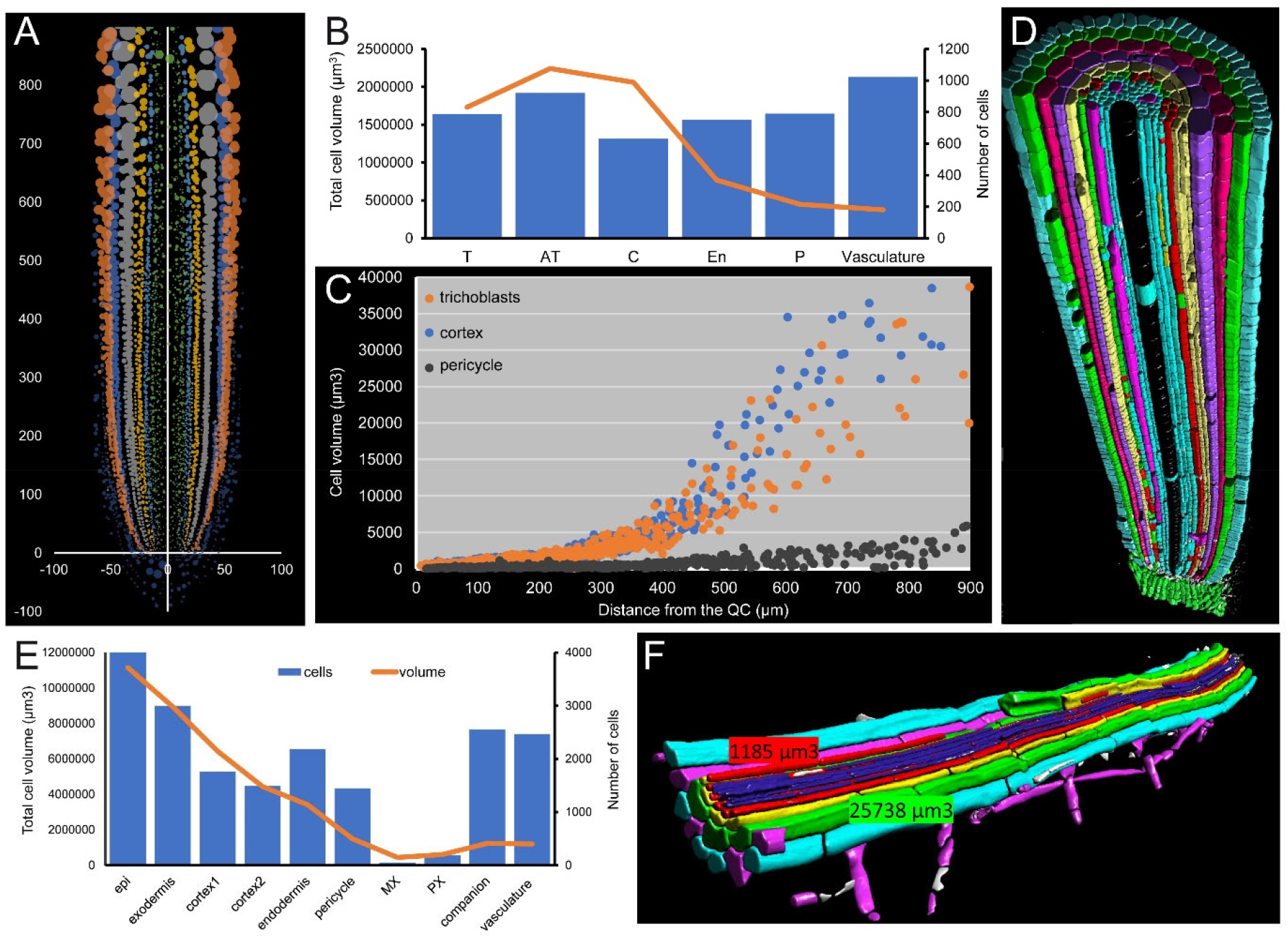
(A) Unrolled root as distance from the main axes. (B) Cell number and cell volume in the first 1 mm from the QC. Bars: cell number; lines: cumulative volume. (C) Cell volume distribution along the axis in the trichoblast, cortex and pericycle. (D) A 3D render of *Setaria italica* root after segmentation. (E) Cell number and cell volume in first 1 mm from the QC. Bars: cell number; lines: cumulative volume. (F) A 3D render of the mature part of the Arabidopsis PR after segmentation. Numbers mean cell volume in the cortex and pericycle cells.

Epidermis is the outer layer of the root and includes two different cell types, trichoblast (T) from which the root hairs are produced, and atrichoblasts (AT) that are radially arranged on a highly constant pattern in the mature Arabidopsis PR: 8 T cell files regularly interspersed between 16 AT cell files (Figure 8A; Guimil and Dunand 2006). Generally, eight epidermis initials directly contact the QC cells in the stem cell region and are the origin of all 24 epidermis files by tangential divisions occurring at variable spatial coordinates (see > in Figure 8B). For a deeper analysis of each tissue, we draw “unrolled root graphs” containing all cells of a given cell type as regards their radial location (x) and distance from the QC (y), which might accommodate other parameters as well, such as cell or nuclei volume (Figure 8B). Unrolled root graphs from same or different samples might be combined to create cellular density maps that allow the detailed study of cellular interactions at the tissue level (Figure 8C). Module IV analysis now allows the detailed mapping of these formative divisions in different genetic backgrounds and in response to different environments (Supplemental Figure S5).

**Figure 8.**
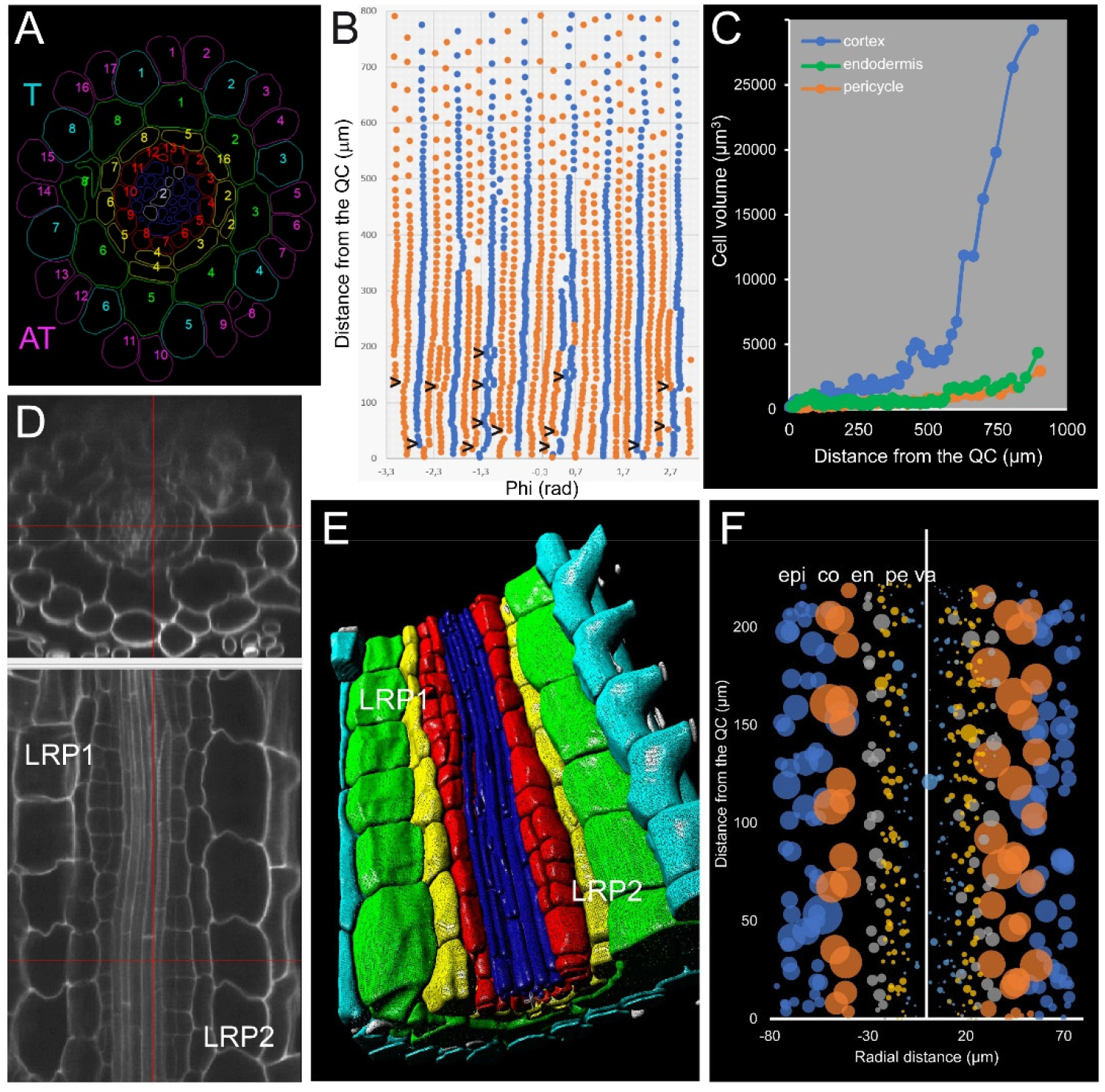
Study of cellular interactions at the tissue level. (A) Virtual cross-section of the PR at the transition zone with the cell files numbers in each cell type; T: trichoblasts, AT: atrichoblasts. (B) 2D graph on an “unrolled epidermis” (blue: T; orange: AT); tangential cell file divergence shown by >. (C) Evaluation of the cell volume in single cell files of cortex, endodermis and pericycle. (D-F) Auxin-induced (100 nM NAA, 18 h) lateral root primordia (LRP). (D) Original 3D scan indicating the position of two LRP (stage II). (E) 3D render after segmentation. (F) Radially unrolled root with cell volume as radius of the circle; epi: epidermis, Co: cortex, en: endodermis, pe: pericycle, va: vasculature.

In addition, we apply module IV analysis to study the effect of exogenous auxin on the morphology of cells at the elongation zone of the PR, confirming the localized and temporal activation of some pericycle cells after a short incubation with 100 nM NAA (Figure 8D-F).

## DISCUSSION

Plant productivity relies on the correct functioning of the different organs with specialized functions within the plant body, and on their interactions with biotic and abiotic factors. Plant organs consist of hundreds or thousands of cells from different tissues, each cell having a determinate gene and protein profile and a fixed spatial allocation and that contribute with specific functions and cell-to-cell interactions. In addition, the shape and size of all plant organs is determined by the contribution of their individual cells through organized cell division and cell expansion patterns. Hence, a comprehensive description of individual cell behavior on a growing organ is crucial for expand our understanding of plant development. Recent advances in tissue labeling, microscopy, and bioimage analysis seize an opportunity to build detailed three-dimensional (3D) representations of plant organs, including morphological, genetic and functional information from each cell.

Roots are key factors in substrate anchoring and nutrient and water uptake, and their simple morphology and the development of tissue-specific fluorescent markers has allowed obtaining gene expression profiles of different root tissues in plant model species such as Arabidopsis (Brady et al. 2007) and tomato (Kajala et al. 2021). This approach relies on the use of molecular markers, which are not directly available in non-model plants, and require protoplast isolation that is known to alter cellular interactions and might affect the expression of some genes. Lowering of next-generation sequencing (NGS) and bioinformatics costs has allowed the use of single-cell RNA sequencing techniques (Jaitin et al. 2014) to characterize the spatiotemporal developmental trajectories of all cells within a root (Zhang et al., 2019; 2021; Shahan et. al. 2021).

Despite these extraordinary advances, many plant biology laboratories lack the appropriate expertise and/or funding to apply the state-of-the-art approaches for quantitative description of root features at cellular resolution. To close this gap, we described here a dedicated pipeline to perform a three-dimensional (3D) analysis of root features at single cell resolution, including root asymmetry, lateral root analysis, xylem and phloem structure, cell cycle kinetics, and chromatin determination. Our pipeline has been organized along four phenotyping modules which require minimal expertise and/or equipment.

Module I was aimed for nuclei and chromatin analysis, and might be used to build chromatin accessibility maps when combined with ATAC-seq and laser-capture microdissection (Chen et al. 2021). This module also allows studying cell differentiation status, as chromatin organization is tightly linked to gene expression regulation and to the differentiation status of the cell. We found that chromatin status in the proliferation zone of the RAM in Arabidopsis roots varied between neighboring cells, even within the same tissue layer. On the other hand, differences in chromatin regularity in the differentiation zone of the PR between Arabidopsis and alfalfa, might account for enhanced cell division during nodule production in legumes.

Module II includes cell cycle analysis aimed for a functional study of root zonation. Several authors defined the meristematic region of the PR in Arabidopsis based on relative cortex cell elongation, without considering other tissues or the actual proliferation activity of the cortex (Dello Ioio et al. 2007; Salvi et al. 2020). Despite the practical value of this definition, module II application allowed us to precisely map cell division, cell endoreduplication and cell elongation in every tissue in every cell (Lavrekha et al., 2017), allowing the building of integrative 3D models of the RAM zonation in species with complex root architecture such as alfalfa, tomato of foxtail millet. Additionally, our protocol will be useful to study processes, such as LR initiation and nodule/pseudonodule induction, occurring in the differentiation zone of the root. Our results also indicate that, shortly upon auxin application, DNA replication occurred not only in the pericycle but in the neighboring procambium of the DZ of the root, while endodermis and cortex cells of this region were unresponsive. By applying module I and II to known auxin mutants and by applying different treatments (i.e. auxin biosynthesis/response inhibitors), the missing link between auxin function and cell cycle regulation could be finally untangled.

Combination of modules I and II will allow to investigate the relationship of cell cycle stages and chromatin organization in each cell, hence comparisons between different tissues and/or regions along the longitudinal root axis will be possible. In addition, comparison between PR, LRs and nodule formation might reveal conserved features of proliferative plant tissues.

Module III includes the detailed investigation of protein localization, histone modifications, and protein complex formation. There are two main methods for the detection of protein localization: fluorescent recombinant protein and immunolocalization. The first one has several disadvantages, like the requirement of gene cloning, protein fusion construction, and plant transformation. Moreover, introducing a fluorescent protein into mutant genotypes requires crossing and it is highly time-consuming. The list of available antibodies with cross-reactivity in other plant species has increased considerably (Oh et al. 2020). As extra benefits, the possibility of colocalization, triple labeling, and protein complex investigation (Pasternak et al., 2018) made this module very attractive for other researchers.

Finally, module IV studies cell geometry features, considered those as a result of epigenetic, gene expression and cell cycle regulation. Size and shape of a given plant organ is ultimately determined by the contribution of its constituent cells through precise division and expansion patterns on a 3D space. As cell size differs drastically across plant organs and cell types, growth coordination at the tissue and organ levels are required (Sablowski 2016). To understand the phenotype of a complex plant organ, such as the root, information about geometry and spatial coordinates of all its cells needs to be obtained. For example, root waving and gravitropic response is causally related to unequal cell elongation in the outer cell layers of the epidermis and cortex (Thompson and Holbrook 2004). By applying module IV to the study of roots of several plant species (Arabidopsis, tobacco, tomato, alfalfa and foxtail millet), we will be able to determine the conserved features of the RAM structure, such as the stem cell niche region and the extent of formative regions in the proximal meristem, among others.

We believe our phenotyping pipeline will allow researchers working on the root biology field to enlarge their Cell Biology toolset, and we do not exclude that other plant researchers working in shoot apical meristem and leaf initiation or *de novo* regeneration will apply our method in their investigations.

## ACKNOWLEDGEMENTS

We thank María José Ñíguez-Gómez for her expert technical assistance and Eduardo Larriba (Universidad Miguel Hernández de Elche, Spain) for useful suggestions. Special thanks to Franck Ditengou, Katja Rapp, and Klaus Palme for fruitful discussions, Thorsten Falk for software modification, and Roland Nitschke with the staff of the Life Imaging Center (LIC) in the Center for Biological Systems Analysis (ZBSA) of the Albert-Ludwigs-University, Freiburg for help with their confocal microscopy resources, and the excellent support in image recording.

## ADDITIONAL INFORMATION

### Authors’ contributions

TP and JMPP designed the research. TP performed the research and analyzed the data. TP and JMPP wrote the manuscript.

### Data Availability Statement

All data that support the findings of this study are available from the corresponding authors upon reasonable request.

